# Copresent microbiome and short-chain fatty acids profiles of plant biomass utilization in rumen simulation technique system

**DOI:** 10.1101/2022.06.22.497131

**Authors:** Tao Shi, Tingting Zhang, Xihong Wang, Xiangnan Wang, Weijun Shen, Xi Guo, Yuqin Liu, Zongjun Li, Yu Jiang

## Abstract

Rumen contents have considerable utility in converting plant biomass to short- chain fatty acids (SCFAs) when used as inoculum for *in vitro* fermentation. To better understand the microbial communities and their functions when *in vitro* ruminal fermentation, the microbiome and SCFAs production were investigated using rumen simulation technique (RUSITEC) system which was inoculated/co- inoculated with rumen contents from goat and cow. This study reconstructed 1677 microbial metagenome-assembled genomes (MAGs) from metagenomic sequencing. The copresent microbiome containing 298 MAGs were found in metagenomic data of these contents and previous ruminal representative samples. These copresent MAGs were overrepresented in decomposing various substrates, especially pectin and xylan. Additionally, the SCFAs productions in RUSITEC were linked with copresent MAGs. Copresent MAGs obtained from this study shows promise to point out the direction for further research on *in vitro* ruminal fermentation, and enables a better understanding of rumen microbiotal structures and functions under *in vitro* condition.

**Graphical Abstract:** 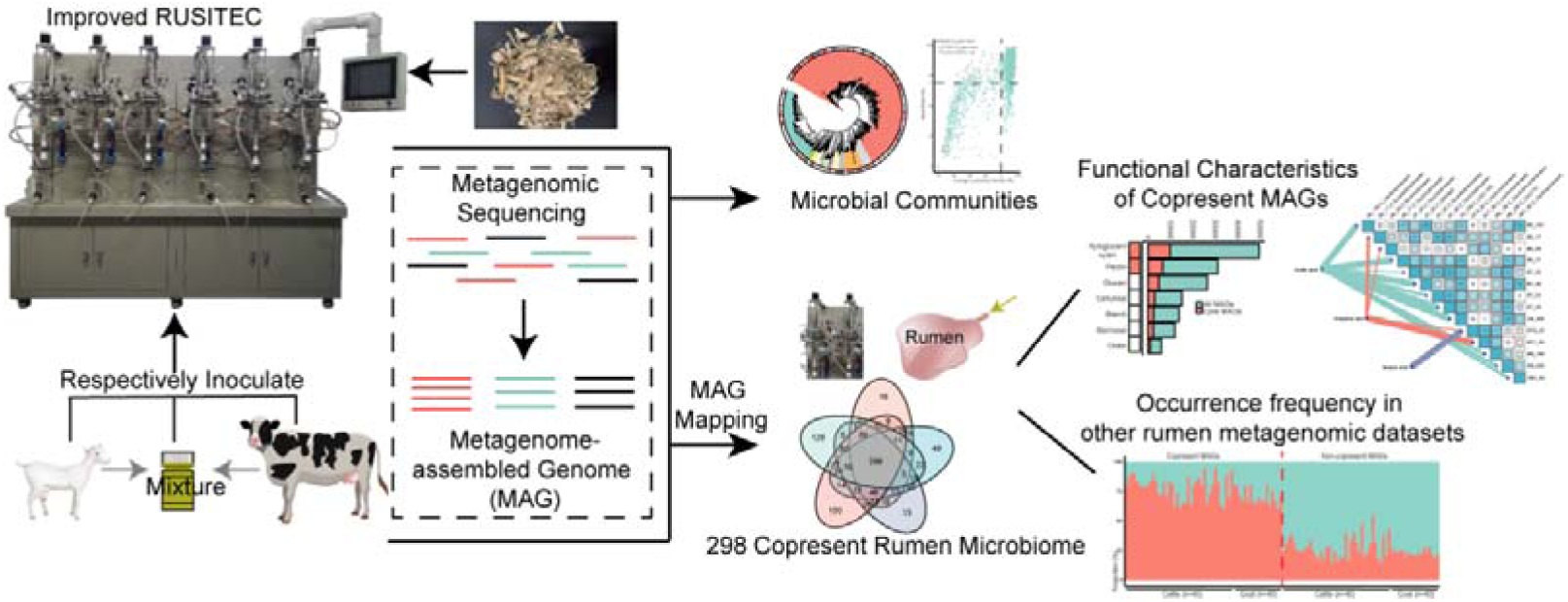

## 1. Introduction

The rumen is a large important part of ruminant digestive system and it is known as natural fermenter because of rumen microorganism’s abilities to efficiently degrading of straw biomass upon large amount of straw biomass intake in ruminants. The potential abilities of rumen microbiomes to degrade lignocellulose are constantly being investigated ^1–2^. Rumen contents of ruminants like cattle and bison have been discovered to continue producing fatty acids and methane during *in vitro* anaerobic fermentation ^3^. A variety of rumen simulation technique systems (RUSITEC) for plant biomass conversion have been developed ^2^. The RUSITEC is an exclusive anaerobic fermentation model that uses rumen contents as the starting inoculum to convert straw biomass to diverse compounds *in vitro*, such as short-chain fatty acids (SCFAs)^4^. Furthermore, the RUSITEC system provided a controlled and stable setting for further experimental modifications of rumen microbial communities without host. Inoculating RUSITEC with synergism of cow and bison rumen contents, for example, resulted in an increase in SCFA production when compared to those singly inoculated with cattle rumen contents ^5^. Rumen microbial communities have the potential to be used for producing high-value products *in vitro* and using a combination of microflora from rumen and other habitats as initial co-inoculum for anaerobic fermentation is something worth looking into. On the other hand, in the process of *in vitro* ruminal fermentation, a clear microbial structure and function are important to improve our understanding of the mechanism of rumen microflora transforming substrates into SCFAs. In a prior research, the 30 most frequent bacterial genera were found in over 90% of rumen and foregut samples from 742 individual animals, indicating that *Bacteroidea* and *Firmicutes* are the most prevalent bacterial phyla in the rumen^6^. While the resolution of these shared microbiomes was classified at higher taxonomic levels, such as genera, this restricts our future investigation of their roles ^7^. The core microbiome of *in vitro* ruminal fermentation and the mechanisms of these microorganisms digesting plant polymers are still unclear, and determining them is the key to use rumen microorganisms to digest plant biomass under *in vitro* anaerobic fermentation conditions.

The construction of metagenome-assembled genomes (MAGs) via metagenomic binning presents a better approach to more precisely expose the microbial population and its activities ^8^. Metagenomic binning was used to reconstruct the strain-level rumen MAGs from ruminant, and the carbohydrate enzymes contained in rumen MAGs have also been explored ^9^. Additionally, using metagenomic binning to investigate the fine-resolution microbial community of anaerobic digestion system has gained more attention recently ^10^. The present study aims to find high-resolution copresent MAGs across host species in natural and *in vitro* ruminal fermentation, and elucidate the abilities of these MAGs degrading plant biomass in improved RUSITEC. The rumen contents of dairy goats, cows and co-inoculum were used as inocula of RUSITEC; subsequently, the SCFAs production was detected as an indicator to evaluate the fermentation efficiency of RUSITEC. Individual taxa’s strain-level microbial genomes were reconstructed using metagenomic sequencing and binning. Meanwhile, the copresent rumen MAGs was identified, and the relationship between these MAGs and SCFAs production was established. This work improved the understanding of the microbial structure in *in vitro* ruminal fermentation and suggested that follow-up research on *in vitro* ruminal fermentation may focus on these copresent MAGs and their functional characteristics to accelerate the application of rumen microbial community.

## 2. Materials and methods

### 2.1 Experimental design and inoculation

Three kinds of rumen contents (from dairy goats, dairy cow, and their mixture) were inoculated in the RUSITEC with four repetitions. The rumen contents were collected 2 h after feeding by permanent rumen fistulas. The rumen collection procedure was approved by the Institutional Animal Care and Use Committee of Northwest A&F University. To standardize the biomass of samples prior to inoculating the RUSITEC, the total DNA concentrations of initial inocula were utilized as the standard for total biomass in each sample ^3^. Precipitate were processed to extract total DNA using the FastDNA^®^ SPIN Kit for Soil (MP Biomedicals, USA) according to the manufacturer’s instructions. However, total DNA concentration was detected with NanoDrop 2000 (ThermoFisher, USA). The inocula were diluted in proportion according to the concentration of DNA using the McDougall buffer solution ^11^.

In RUSITEC, two rounds of *in vitro* fermentation were performed with six reactors. Each round experiment included two cow rumen contents (cow group), two goat rumen contents (goat group), and two mixed contents of cow and goat rumen (co-inoculum group). A modification was made based on the previously described method of diluting inoculum ^12^, which takes the total biomass as a consideration. Two rounds of cow rumen inocula (DNA concentration: 28.93 ± 2.91 μg/mL; 23.65 ± 0.27 μg/mL) were diluted 1.18 and 1.28-fold respectively with McDougall buffer solution, while the goat rumen inocula (DNA concentration: 24.48 ± 0.15 μg/mL; 18.42 ± 0.69 μg/mL) were diluted 0.84 and 0.77-fold respectively. For co-inoculum groups, the proportions of two rounds of experiments were V_cow_: V_goat_: V_McDougall buffer_ = 1:1.18:2.18 and 1:1.28:2.28 respectively. Finally, the total volume in the reactor was controlled to 1 L, and the total DNA concentration was 13.25 μg/mL. Four initial rumen samples from two cows and two goats, and 12 final samples in anaerobic reactors were collected for metagenomic shotgun sequencing analysis.

### 2.2 RUSITEC Experiment

The night before inoculation, reactors of RUSITEC were preheated and maintained at 39 ± 0.5°C. Nitrogen gas was infused into the fermenters to maintain an anaerobic environment. As fermentation progresses, the McDougall buffer solution was used to maintain the pH, buffer dilution rate was set to 1.5%. The dried whole-plant corn and concentrate (60% corn grain, 12% wheat bran, 20% soybean meal, 3% rapeseed meal, 1.5% calcium, 1.5% salt, and 2% premix) were used as substrate for fermentation, crush the dried feed and sieved it with 40 mesh screen. Each RUSITEC fermenter was fed twice daily at a 12-hour interval, with a concentrate-to-roughage ratio of 10 g: 10 g each time. Furthermore, the microbial communities of various inocula were modified in RUSITEC for 7 days. The air pressure, pH, and temperature in the fermenters were monitored in real time during the experiment, and the stirrer ran constantly to maintain a consistent environment for *in vitro* ruminal fermentation.

### 2.3 Sample collection and processing

The contents of fermenters were collected at the sample port after a 7-day stabilization period. Sample processing and chemical analysis were performed as previously described ^3^. Supernatants from samples were separated for testing SCFAs concentrations using Agilent 7820A Gas Chromatograph System (Agilent Technologies Inc., USA); pellets from samples were used for metagenomic shotgun sequencing. The DNA extraction process of pellets was performed refer to the manual of MagPure Stool DNA KF kit (Magen Biotechnology Co., Ltd., China). After DNA quality control and fragmented, the selected DNA fragments were repaired, and then ligated with indexed adaptor. Finally, processed samples were sequenced by DNBSEQ Platform (MGI Tech Co., Ltd., China) after two rounds of PCR.

### 2.4 De novo metagenome assembly, binning and genome annotation

The metagenomic shotgun sequencing data were processed as previously described ^13^, including metagenomic assembly, binning, dereplication, genome annotation, and Carbohydrate-Active enZYmes (CAZymes) domain, etc. In addition, thresholds of completeness ≥ 50% and contamination ≤ 10% were used to filter MAGs. Phylogenetic trees of MAGs were reconstructed using IQ-TREE (v 1.6.9) ^14^; the visualization and annotation of phylogenetic trees were carried out in iTOL (v 6.5.1) (https://itol.embl.de/). The clusters of KEGG pathways were annotated using KofamKOALA (v 1.3.0) ^15^. Estimated genome size of MAG was corrected as described in previous study using completeness and contamination ^16^. In addition, the obtained MAGs were compared to the rumen MAGs set collected from previous studies (n = 22506) ^9, 17–20^ and the GTDB representative genomes (n = 31910) using fastANI (v 1.1) ^21^.

### 2.5 Regrouping MAGs into copresent and non-copresent MAGs

Mapping method was used for regrouping MAGs. First, bwa -a bwtsw (v 0.7.13)^22^ was used to build an index based on all MAGs. Then, the metagenomic data of each sample were mapped back to MAGs index using bwa -mem. And MetaSNV (v 1.0.3) ^23^ was used to calculate the average coverage for each MAGs in BAM files. As the cut-off of MAG’s presence/absence, average coverage was adjusted using data size of each sample, and copresent MAGs were defined as the MAGs presented (average coverage > 1) ^20^ in every sample of cow, goat, co-inoculum group, and initial inocula. The rest of MAGs couldn’t meet the threshold in all samples were defined as non-copresent MAGs. To determine whether our MAGs could present in other rumen metagenomic data, 40 randomly selected samples in previous study ^9^ and 18 goat rumen metagenomic data under different dietary feed ^24^ were downloaded from their sources and mapped to the 1677 MAGs, the average coverage and percentage coverage of copresent MAGs and non-copresent MAGs were extracted from the obtained mapped results for subsequent analysis. As mentioned above, average coverage > 1 and the means of percentage coverage in the results of goat and cow samples mapped to 1677 MAGs were used as the cut-off of presence/absence in samples.

### 2.6 The interaction between SCFAs production and copresent MAGs abundance

In order to find the genome-level evidence of producing SCFAs in copresent MAGs, EC number of enzyme were selected according to prior study ^25^. Next, Pearson correlation analyses were performed on SCFAs concentrations and the abundance of copresent MAGs using R package “hmisc”. The final results showed the positive correlation (r > 0.75; FDR adjusted *p*-value < 0.05) between SCFAs concentrations and copresent MAGs with the genome-level evidence of producing SCFAs, as well as the correlation between these MAGs.

### 2.7 Statistical analysis

One-way ANOVA and turkey pairwise comparisons were used to examine the significance of differences in SCFA concentrations between cow, goat, and co- inoculum groups. Fisher’s exact test was employed to test the enrichment of the CAZyme/KEGG pathway/family-level count in copresent MAGs *vs*. all MAGs. Corrections for *p* values were applied whenever stated using R package “p.adjust”, FDR adjusted *p*-value was also expressed as Q-value.

## 3. Results and discussion

### 3.1 Effect of inoculum sources on short-chain fatty acids production in RUSITEC

SCFAs production was evaluated in RUSITEC which was inoculated with different microbial inocula. Acetic acid, the primary component of SCFAs, was discovered in goat, cow, and co-inoculum groups at a concentration of 54-65%; propionic acid and butyric acid represented for 17-22% and 13-19% of total SCFAs concentration in these groups, respectively. In contrast, the low-level proportion of valeric acid, isovaleric acid and isobutyric acid were detected in three groups (Figure 1). These are the common ratios of SCFAs during ruminal *in vitro* and *in vivo* fermentation ^26–27^. The differences of specific SCFA concentration were further analyzed. It was found that the concentration of acetic acid was the highest (12138.28± 467.85 mg/L) in cow group, while the lowest concentration in goat group (9751.83± 1092.87 mg/L) (*p* < 0.05). The highest and lowest concentrations of propionic acid were found in the cow (5005.90± 499.86 mg/L) and goat (3693.0± 465.28 mg/L) groups, respectively, following the same pattern as acetic acid (*p* < 0.05). For the butyric acid, concentrations in cow group (4854.69± 915.04 mg/L) were also higher than that in goat group (3275.70± 340.47 mg/L) (*p* < 0.05), and there was no significant difference in the concentration of isobutyric acid between three groups. This result was supported by previous studies, which also found that rumen contents from cows had higher fermentation and degradability than dairy goat, both under *in vivo* and *in vitro* conditions ^28^. For lower concentrations of valeric acid and isovaleric acid, the concentrations were significantly higher in cow and co- inoculum group compared to goat group (*p* < 0.01) (Figure 1). Overall, in RUSITEC system, different inocula resulted in differences of specific SCFA concentration between groups, and reactors inoculated with cow rumen contents had the highest capacity in producing SCFAs, followed by the co- inoculum group. In addition, co-inoculated with cow rumen content in proportion, the abilities of producing special SCFAs in RUSITEC were improved compared to individually inoculated with goat group.

**Figure 1.**
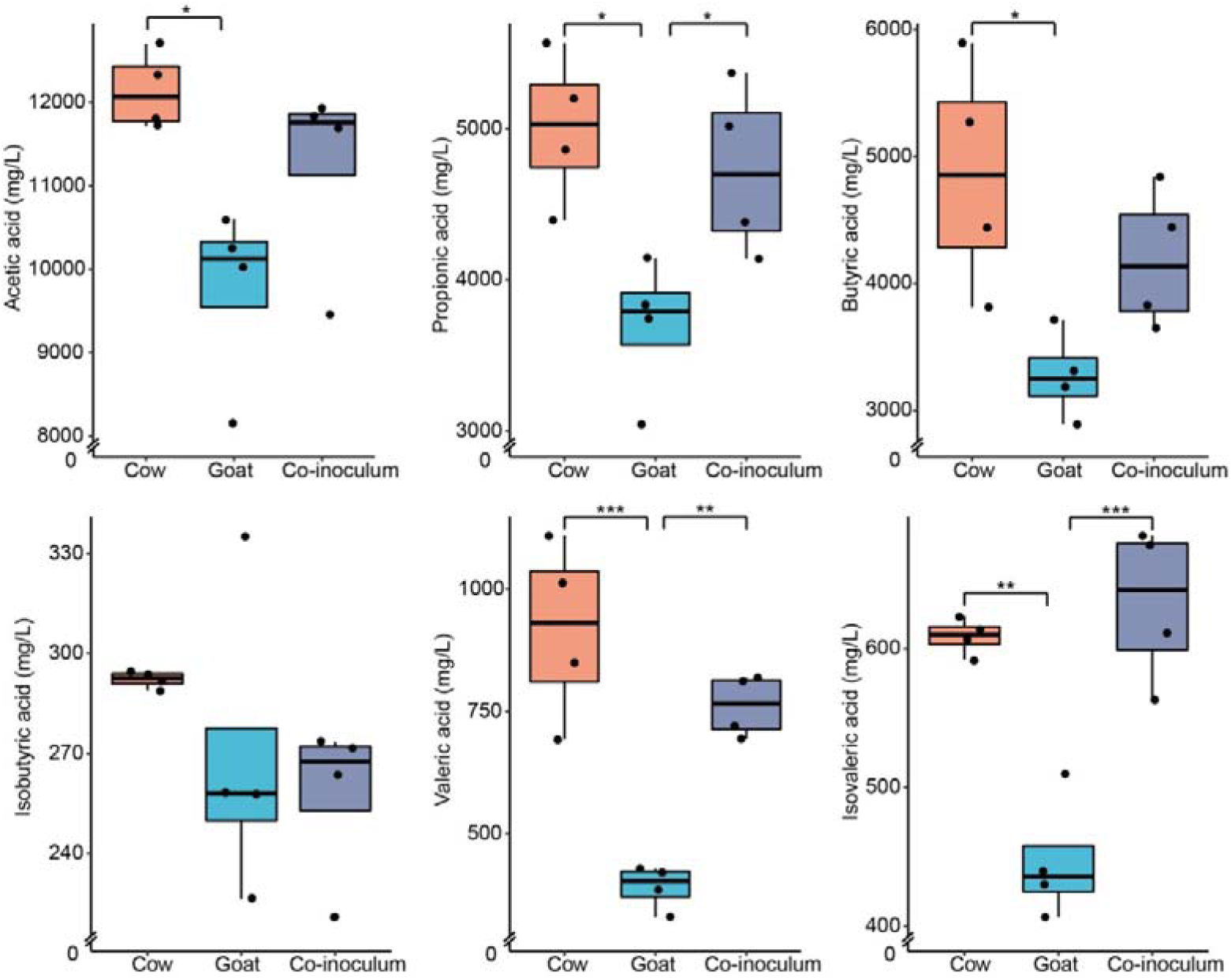
The differences of SCFAs production in the RUSITEC between three groups. Cow and goat, respectively represent RUSITEC inoculated with rumen contents from cow and goat, co-inoculum represents RUSITEC inoculated with the mixtures of biomass-normalized rumen contents from cow and goat (* Q-value < 0.05, ** < 0.01, *** < 0.001).

The potential of ruminal microbiomes to convert plant biomass to various high- value compounds *in vitro* was developed through the research of *in vitro* ruminal fermentation ^2, 26, 29^. However, the control of fermentation environment *in vitro* is one of the main factors leading to various results. Otherwise, it was confirmed that the rumen fermentation efficiency of cows was higher than that of goats, there were still follow-up studies that obtained the opposite results and it was difficult to give a reasonable explanation ^30^. It could be seen that a standardized *in vitro* ruminal fermentation system and experimental process are urgently needed to develop rumen microflora as effective resources for degrading cellulose. The RUSITEC was partially improved as the fermentative system used in this study, and it is now equipped with a completely automated control system that allows it to inoculate 1 L of rumen inoculum and sustain long-term and stable anaerobic fermentation conditions for *in vitro* ruminal fermentation.

Importantly, anaerobic fermentation carried out at agitated and non-agitated conditions would make great differences in microbial communities and SCFAs production ^31^. To ensure uniform fermentation in the RUSITEC system, the contents of reactors were continually stirred using an electric motor. On the other hand, the biomass of the initial inoculum had been considered as one of the main variables, which was more conducive to explore the different SCFAs productions caused by different rumen inocula under the same total biomass standard ^3^.

For rumen microflora, different ruminants have different characteristics when it comes to decomposing plant biomass ^32^, and mixing two or more types of unique rumen contents will improve the abilities of *in vitro* ruminal fermentation ^5, 33^. In the current study, the co-inoculum group inoculated with biomass- normalized rumen contents from dairy goats and cows was investigated. Increased SCFAs production was also observed in the co-inoculum group compared to the goat group, while the specific ecological process in the co- inoculum group needs to be investigated further. In conclusion, the performance of co-inoculum in RUSITEC provides new insight into the efficiency and microbial communities of *in vitro* ruminal fermentation, as well as a new vision for blending microbial communities from various habitats in the future.

### 3.2 Metagenome assembled genomes were reconstructed from metagenomic data of initial inocula and RUSITEC samples

In metagenomic sequencing data from different RUSITEC samples (n = 12) and initial inocula (n = 4), generated 4.68 billion clean read-pairs in total, and over 1.4 T metagenomic sequencing data. Then, the 1677 genomic bins were approved as MAGs (completeness ≥ 50% and contamination ≤ 10%) by metagenomic binning with 99% average nucleotide identity (ANI). All these MAGs, which were classified into 18 different bacterial phyla and 2 archaeal phyla, were divided into 186 high-quality MAGs and 567 medium-quality MAGs (Figure 2A and Figure S1). Among 1677 MAGs, 79.8% of MAGs were classified into *Firmicutes*_A (59.7%) and *Bacteroidetes* (20.1%). We reconstructed strain- level individual microbial genomes under *in vitro* ruminal fermentation conditions, as predicted, and discovered that MAGs assigned to the *Bacteroidetes* and *Firmicutes* phyla still dominate the RUSITEC microbiome (Figure 2B). Also, the metagenomic bins in RUSITEC means that follow-up research on microbiome under *in vitro* fermentation conditions can now focus on genomic function of individual taxa or the function of a specific microbiota, as well as having important implications for finely understanding the functional and taxonomic characteristics of rumen microorganisms that can survive *in vitro* under anaerobic digestion.

**Figure 2.**
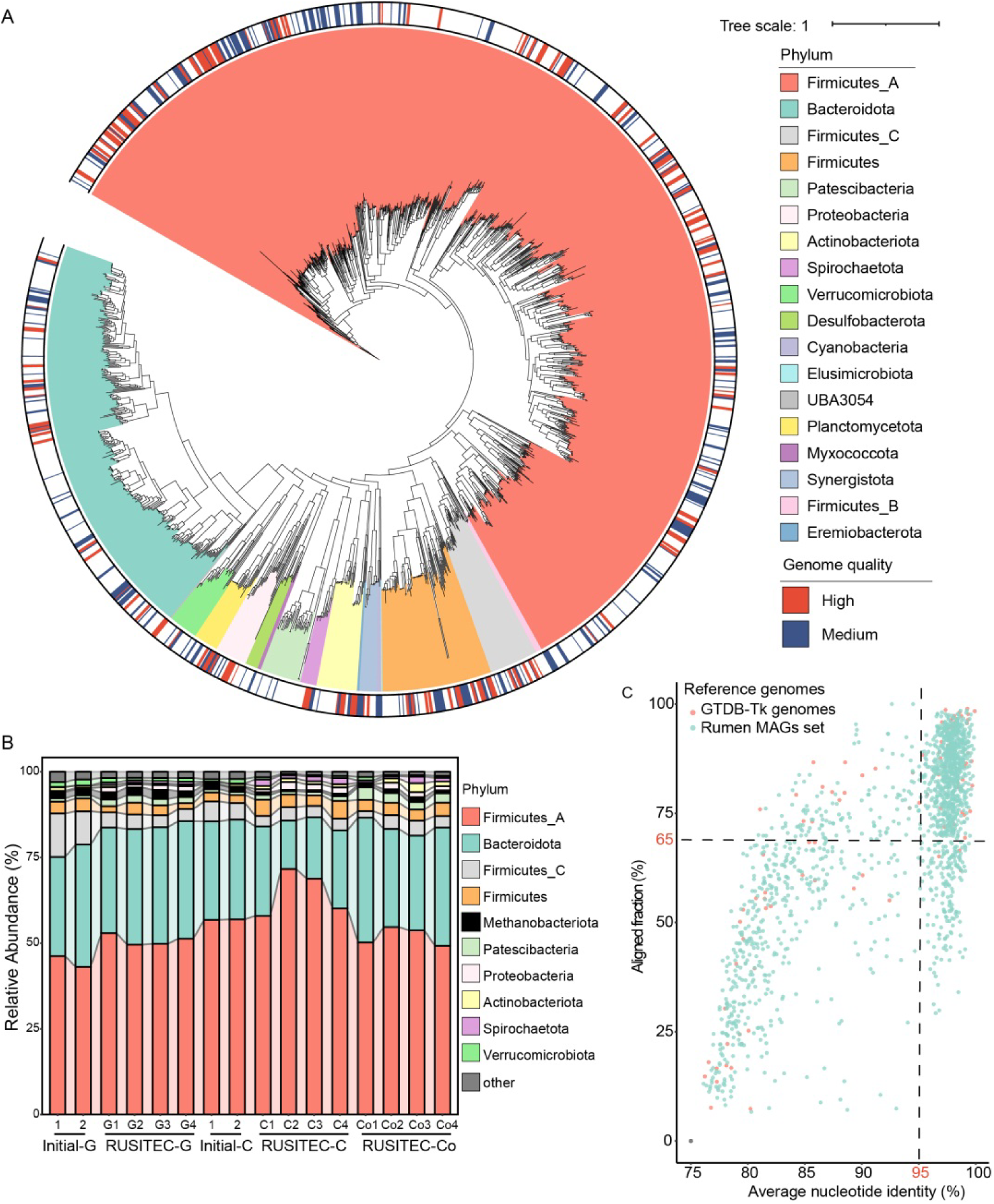
The microbial communities in different samples of RUSITEC and initial inoculum. (A) A maximum likelihood phylogenetic tree of bacterial MAGs. The colored inner ring represents the different phylum of MAGs, colored outer ring corresponds to the genome quality of each MAGs. (B) The phylum-level proportion of MAGs, it shows the the top 10 phyla in abundance. (C) The average nucleotide identity (ANI) and aligned fraction of the 1677 MAGs compared to the rumen MAGs set and the GTDB representative genomes.

For 1677 MAGs, whole-genome ANI were computed with the genomes of GTDB database (n = 31910) ^34^ and the published rumen MAGs set (n = 22506) ^9, 17–20^. It was discovered that 906 of the 1677 MAGs (∼ 54%) were considered as the same species-level MAGs with an ANI > 95% over ≥ 65% of alignment fraction (Figure 2C), and also these MAGs lacked representative in GTDB databases. Among the 906 MAGs met the threshold, only 58 MAGs met an ANI > 95% over ≥ 65% of alignment fraction with GTDB representative genomes. On the contrary, it was observed that 848 MAGs met this threshold with published rumen MAGs set, and 64 of these MAGs had an ANI > 99%. Different from prior research on rumen MAGs ^9, 19^, there were fewer unknown genomes obtained in this study. Whereas, more than 93% of MAGs met the specie-level ANI threshold with rumen MAGs in published studies. This suggesting that the published rumen MAGs were overrepresented in the MAGs set derived from *in vitro* ruminal fermentation, indicating that the ruminal *in vitro* MAGs may share more microbiome with the already published rumen studies.

### 3.3 Copresent rumen microbiome obtained from different habitats

The approach of MAG mapping was utilized to detect the MAGs included in each group according to previous study ^20^. Finally, results showed that 491 MAGs meet the threshold of average 1× coverage in 4 samples of cow group, 648 MAGs in 4 samples of goat group, 621 MAGs in 4 samples of co-inoculum group, 795 MAGs in 2 samples of initial goat inoculum, and 907 MAGs in 2 samples of initial cow inoculum (Figure S2). Next, the presence/absence of MAGs in every sample was used to generate a binary matrix for principal co- ordinates analysis (PCoA). The PCoA results further revealed the high similarity (R = 0.919, *p* < 0.01) within five types of samples (Figure 3A), indicating the composition of microbial structure changed in RUSITEC samples after the stable period compared to the initial inocula. However, there were 298 copresent rumen MAGs in the host’s rumen and RUSITEC contents, as shown in Figure 3B. So far, strain-level microbiome existed stably under *in vivo* and *in vitro* conditions were found. It could improve understanding for rumen microbial abilities on substrate degradation under different conditions.

**Figure 3.**
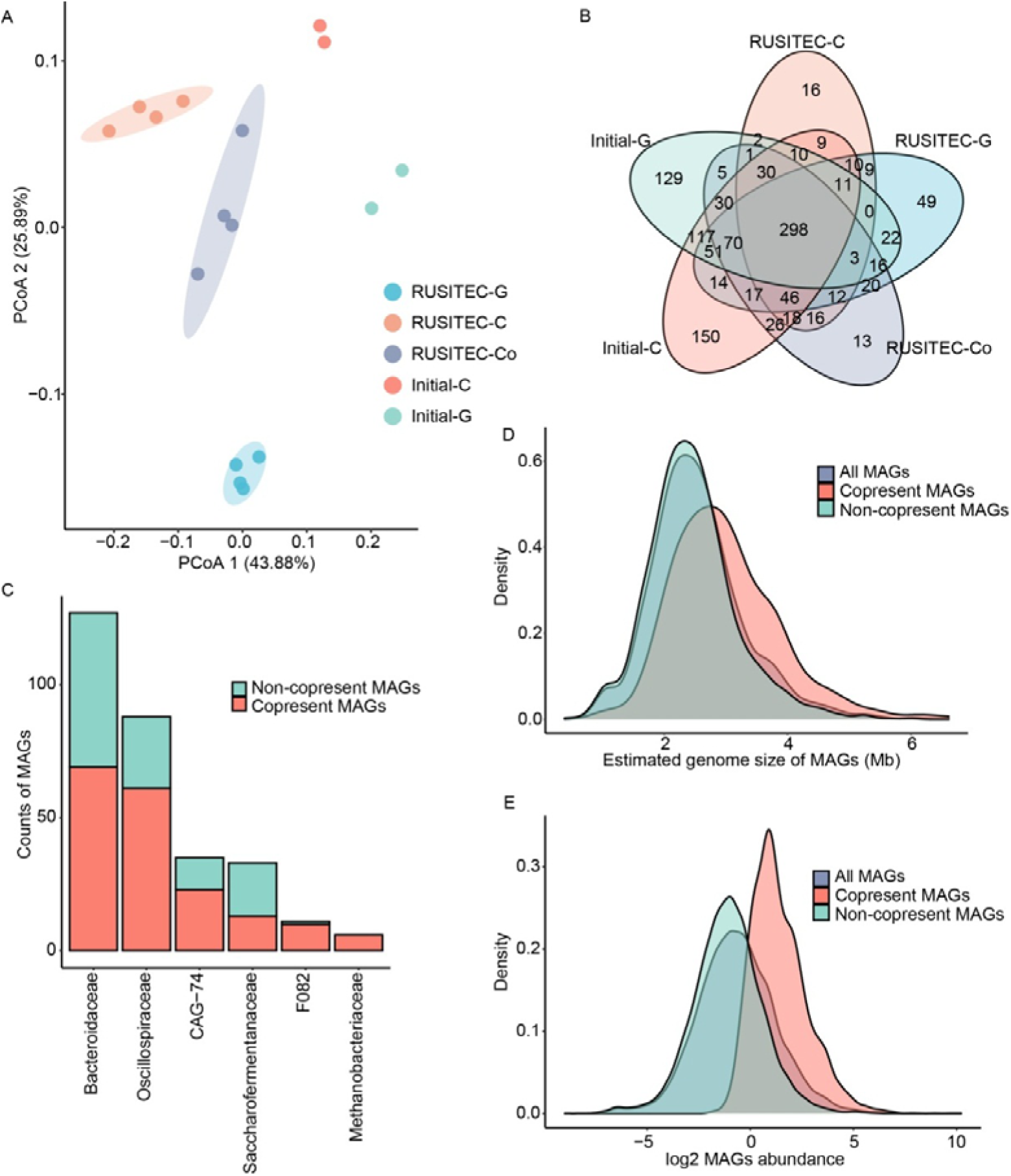
A copresent rumen microbiome consist of 298 MAGs persisted across in different RUSITEC samples and initial inocula. (A) Principal coordinates analysis of abundance of MAGs in each sample based on Bray- Curtis distance. (B) Venn diagram shows copresent rumen MAGs persisted across five types of samples. (C) Histogram shows that bacterial families which are overrepresented in copresent rumen MAGs compare with all MAGs (Fisher’s exact test, Q-value < 0.05). (D-E) Density distribution of the corrected estimated genome size and the log2 abundance of each MAG.

Further analysis demonstrated that these copresent microbiomes were classified into 28 families, and more information about 298 copresent MAGs was shown in Table S1. Figure 3C showed families significantly enriched in the copresent microbiome, including *Bacteroidaceae*, *Oscillospiraceae*, *CAG-74*, *Saccharofermentaceae*, *F082* and *Methanobacteriaceae*. Significantly, six copresent archaeal MAGs, which classified into genus *Methanobrevibacter*, were obtained in copresent rumen microbiome under *in vivo* and *in vitro* conditions in this study. The genus *Methanobrevibacter* has been proved to play a major role on rumen methane production ^35^, elucidating the ecological role of archaea in RUSITEC appears to be further investigate.

Next, we investigated the distribution of estimated genome size and abundance of copresent and non-copresent MAGs in different samples. Furthermore, it was found that larger genomic size and greater abundance were found to be overrepresented in copresent MAGs *vs*. non-copresent or all MAGs (*p* < 0.05). Previous research has revealed that microorganisms with a higher abundance may assume a dominating position in the rumen habitat, and these bacteria have an advantage in resource acquisition over others. On the other side, it has been suggested that a bigger microbial genome size may be associated with a slower growth rate ^36–37^. It was shown that the high abundance and big genome size of rumen microorganisms were two of the factors influencing whether *in vivo* rumen bacteria can persist stably under *in vitro* ruminal fermentation conditions.

To investigate the functional properties of copresent MAGs further, a matrix of KEGG pathway counts was generated for 1677 MAGs, which were then used for further enrichment analysis of copresent MAGs *vs*. all MAGs. Also, we found that the 48 KEGG pathways were substantially enriched (Q-value < 0.05) in the copresent MAGs relative to all MAGs, including amino acid metabolism, carbohydrate metabolism and other pathways related to substrate utilization. While the remaining pathways, including two-component systems, quorum sensing and cell motility, which are critical for bacterial survival, were considerably enriched in copresent rumen MAGs (Figure S3). Furthermore, these overrepresented pathways might be linked to the proper allocation of life history strategies like resource acquisition and stress tolerance ^36^. Overall, the copresent MAGs were discovered for the first time under *in vivo* and *in vitro* fermentation conditions in the cow, goat, and co-inoculum groups, and the genomic and functional characteristics of MAGs were preliminarily explained with finer resolution, advancing our understanding of the copresent MAGs under *in vitro* ruminal fermentation.

### 3.4 Occurrence frequency of copresent rumen MAGs in other rumen metagenomic datasets

In order to evaluate extensibility of 298 copresent MAGs, 58 rumen metagenomic data of cattle and goat were used to generate a datasets for testing ^9, 24^. The average coverage of these copresent MAGs (mean: 10.34X and 4.46X) is higher than that of non-copresent MAGs (mean: 1.93X and 1.37X) in the mapping results of these data mapped to all 1677 MAGs, and the percentage of copresent MAGs (54.54% and 48.58%) and non-copresent MAGs (28.34% and 27.12%) also showed the same trend (Figure 4 A-D). It showed that in the same context of MAG mapping, rumen metagenomic datasets from other studies had the better coverage on copresent MAGs. Then, according to the results of two metagenomic datasets mapped to all 1677 MAGs, selecting depth > 1, and the mean of percentage (cow: 30.06%; goat: 30.93%) as the evidence that a MAGs existed in the corresponding dataset. Figure 4E showed that there were 179-238 copresent MAGs met the threshold in the cattle dataset, accounting for 60% - 95% of copresent MAGs. However, only 14% - 55% (201-765 MAGs) non-copresent MAGs met the threshold. In the goat dataset, despite the changes of dietary conditions, still 60% - 83% of copresent MAGs were found to meet the threshold, and only 18% - 28% of non- copresent MAGs met the threshold. In short, more than 60% of copresent MAGs were detected in every collected metagenomic data, and only more than 14% of non-copresent MAGs were detected in the sampled data. These findings further suggested that there were shared microbiome between the microflora including MAGs recovered from ruminal *in vitro* metagenomic data and other rumen microflora in different geographies and dietary conditions, and the scope of this shared microbiome dataset was narrowed in this study.

**Figure 4.**
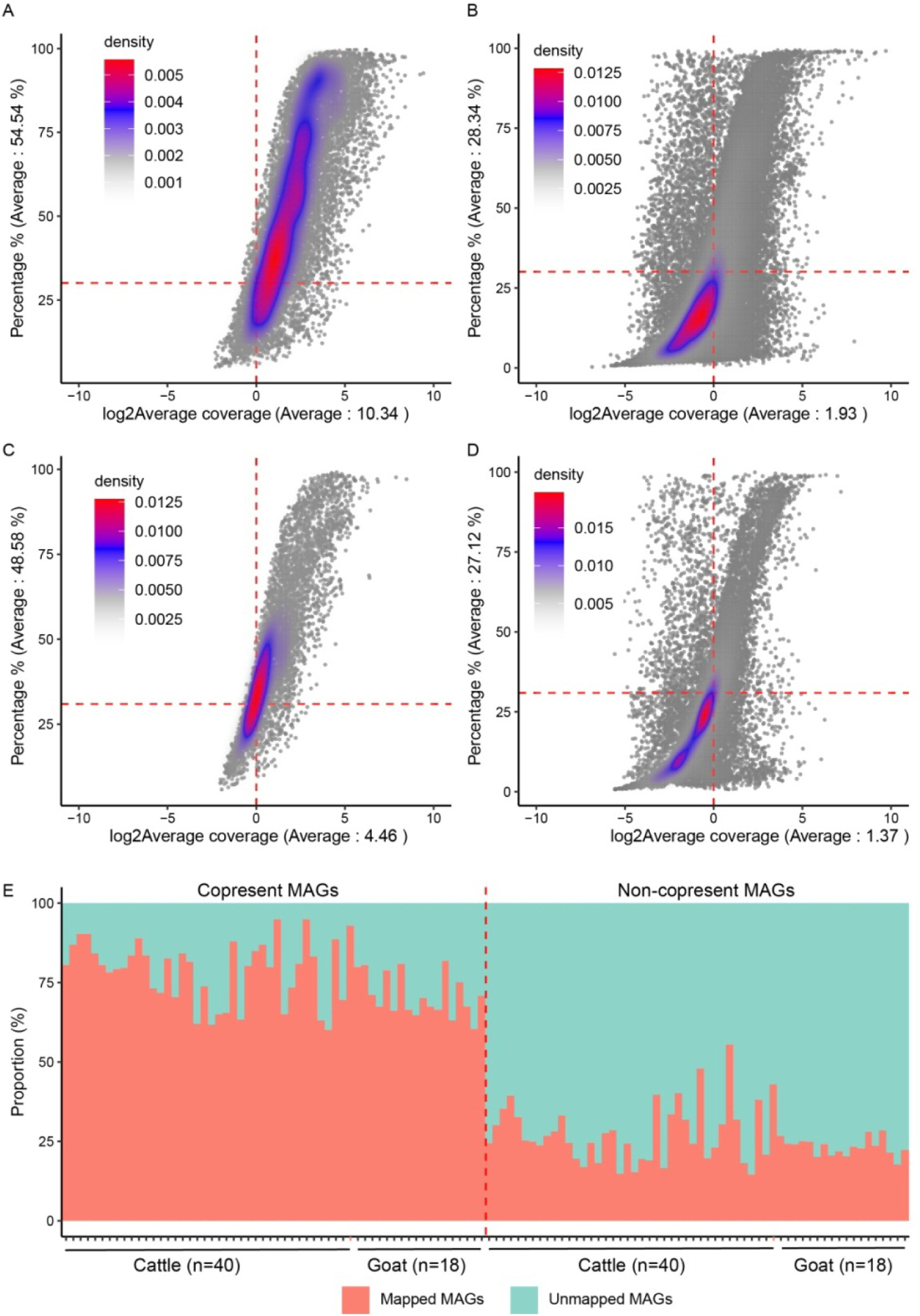
The distribution of average vertical coverage and horizontal coverage obtained from metagenomic dataset mapped to MAGs. (A-B) The mapped result distribution of copresent MAGs and non-copresent MAGs extracted from total mapped result of cattle samples. (C-D) The mapped result distribution of copresent MAGs and non-copresent MAGs extracted from total mapped result of goat samples; log2 average coverage represents vertical coverage, percentage represents horizontal coverage, and the color in the figure represents the density of data distribution. (E) The proportion of mapped MAGs and unmapped MAGs in 298 copresent MAGs (on the left of dash line) or 1379 non-copresent MAGs (on the right of dash line), respectively.

Previously it was observed that geographical and dietary variations were the primary variables influencing the microbial structure of rumen ^6^. The copresent MAGs found in this work, both *in vitro* and *in vivo*, were still found in rumen metagenomic datasets from various geographical locations and dietary circumstances. It was also hypothesized that when additional rumen contents were employed for *in vitro* fermentation, the copresent microbiome would be more likely to survive and play a key role. This finding not only confirmed the presence of copresent MAGs in other datasets, but it also broadened the ranges of copresent MAGs in *in vitro* ruminal fermentation.

### 3.5 Various carbohydrate-active enzymes were overrepresented in copresent rumen microbiome

The 298 copresent rumen MAGs accounted for 17.8% of all MAGs, while the overrepresentation of CAZyme domains in copresent MAGs *vs*. all MAGs were observed in this study. It was found that the polysaccharide lyase (PL) domains contained in the copresent rumen microbiome occupied the highest proportion (34.91%) relative to PL domains contained in all MAGs, glycoside hydrolase (GH) domains in copresent rumen MAGs accounted for 24.15%, and the proportion of carbohydrate-binding module (CBM) (24.39%), S-layer homology module (SLH) (24.51%) and auxiliary activities (AA) (24.02%) domains in copresent rumen MAGs also exceeded 17.8% (Figure 5A). Furthermore, the CAZyme domains enriched in copresent MAGs were investigated as previously described ^38^, found that cellulase-related domains (e.g., GH5, GH51), amylase-related domains (e.g., GH13), and pectin-related domains (e.g., PL1, PL11) were highly (Q-value < 0.05) enriched in copresent rumen MAGs (Figure 5B). Although we have some findings about the overrepresentation of copresent rumen MAGs in specific CAZyme domains, all CAZyme domains were classified into clusters based on their functions on utilizing seven types of plant polymers in order to better understand the advantages of these MAGs in degrading plant biomass. It was found that domains, which can play a role in degrading xylose/xylan and pectin, were enriched in copresent rumen MAGs (Q-value < 0.05) (Figure 5C). It showed the contribution and preference of copresent rumen microbiome in degrading plant polymers. Despite being underrepresented in degrading other plant polymers, it preserved a substantial number of related domains in copresent MAGs.

**Figure 5.**
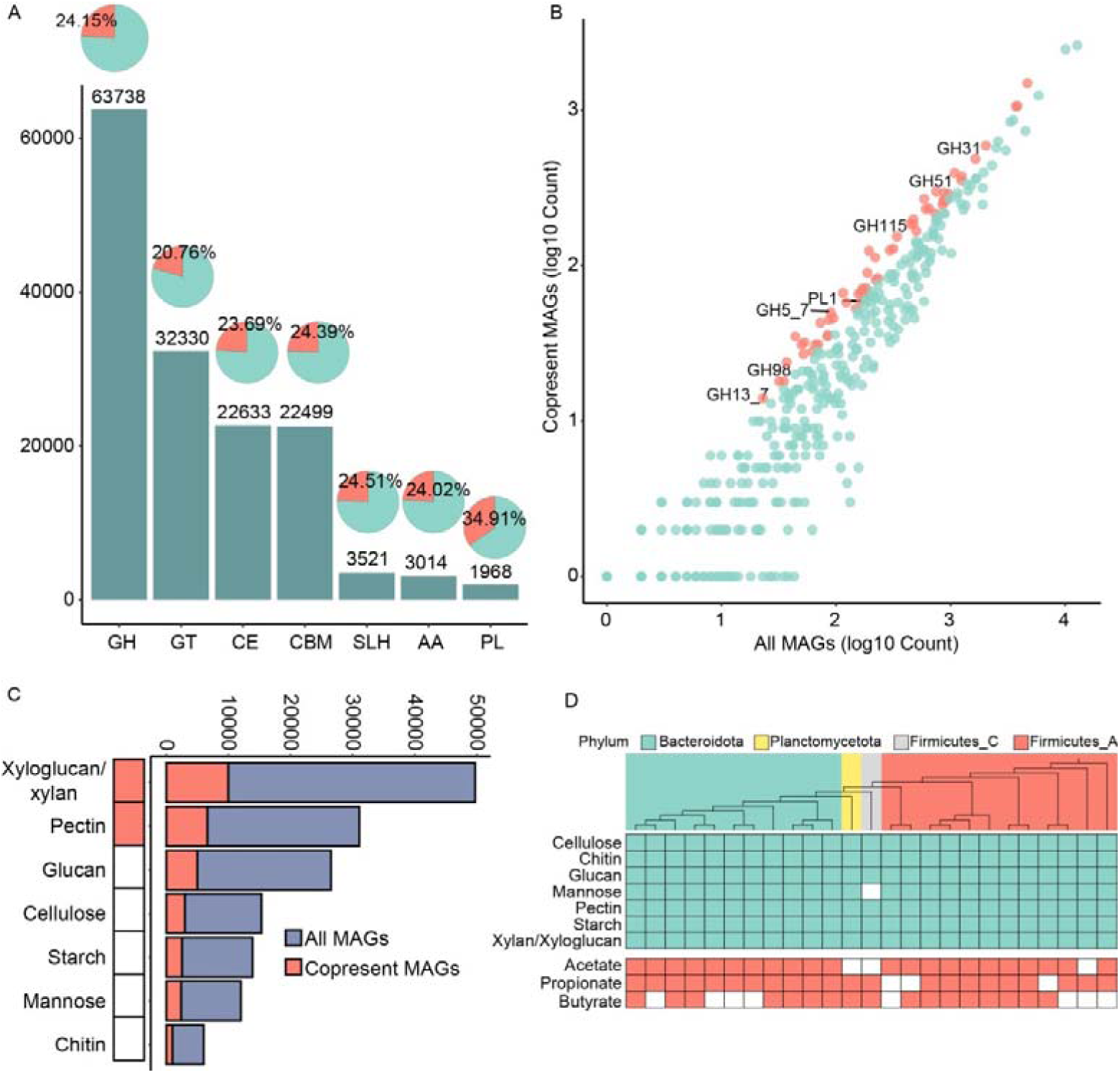
Proportion of carbohydrate-active enzymes contained in copresent rumen MAGs and the abilities of these MAGs in degrading plant polymers. (A) The total counts of CAZymes contained in all MAGs and the proportion occupied by copresent rumen MAGs in special CAZymes (percentage in pie chart). (B) The enrichment of CAZymes domains in copresent rumen MAGs (red points) (Fisher’s exact test, Q-value < 0.05). (C) Overrepresented plant polymers in copresent MAGs compare with all MAGs. The red/green bars are used to represent the total counts of gene in copresent/all MAGs in degrading specific plant polymers. The heatmap display the level of enrichment or depletion based on Fisher’s exact test, colored (significant) cells have Q-value < 0.05. (D) The concerned function of all 25 unique near-complete bacterial genomes in this copresent MAGs.

**Figure 6.**
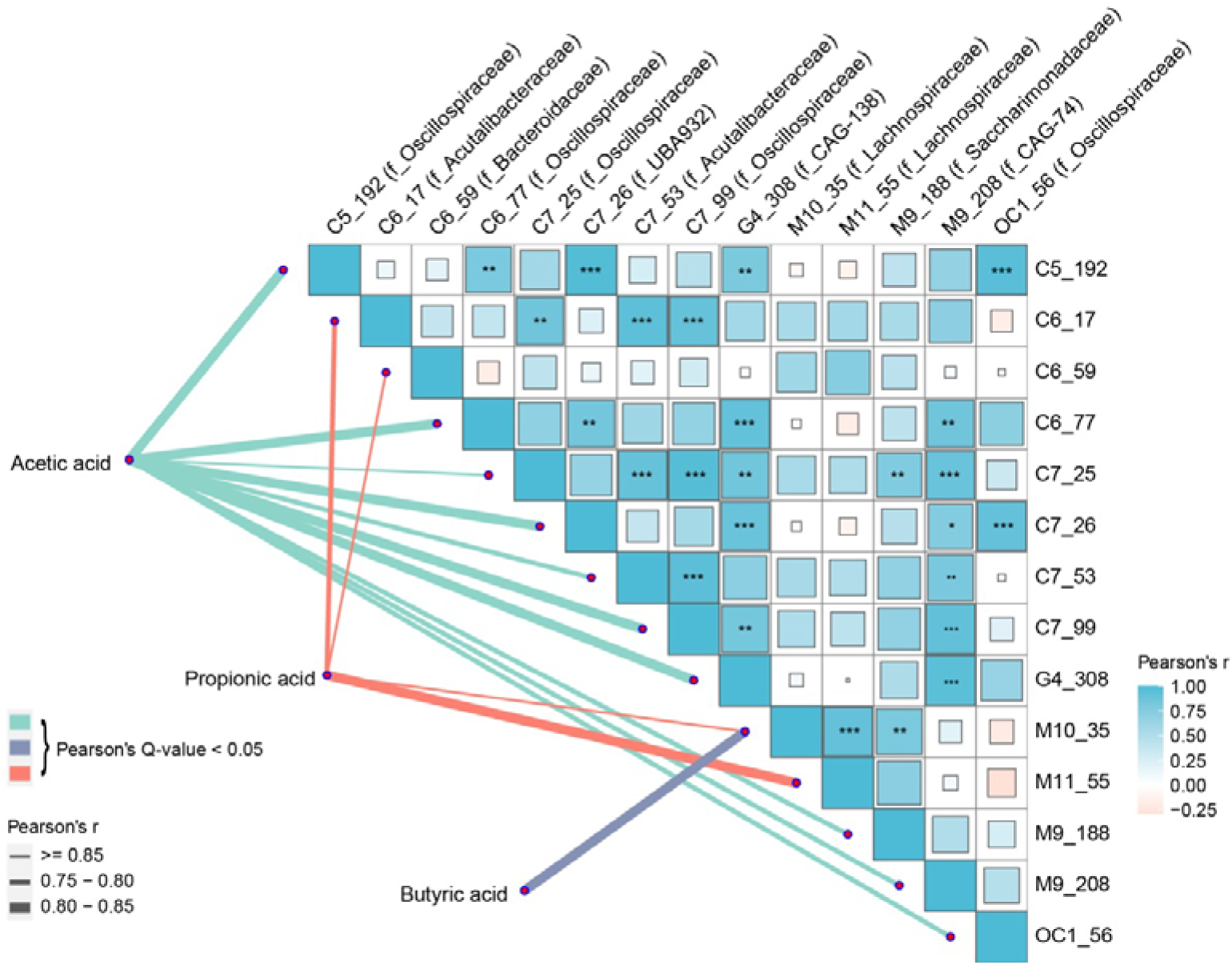
Pearson correlation analysis between abundance of copresent MAGs and SCFAs concentration in RUSITEC. Only the relationships with significant and positive correlation (r > 0.75, Q-value < 0.05) between MAGs and SCFAs concentration are shown in the figure. The lines in the lower left corner represent the correlation between screened copresent MAGs and three types of SCFAs. The heatmap in the upper right corner represents the correlation between these MAGs (*Q-value < 0.05, ** < 0.01, *** < 0.001).

Furthermore, metabolic reconstruction of near-complete MAGs can help to improve the reliability of functional features ^25^. There were 25 near-complete copresent MAGs scattered throughout four phyla, with a completeness > 90% and contamination < 5%. CAZyme domain clusters and important genes of three final products were found in near-complete MAGs, as shown in Figure 5D. Except for one MAGs of *Firmicutes*_C, which had a loss of CAZyme domains in degrading mannose, the other MAGs contained relevant domains for degrading all seven substrates, and one or more abilities of producing three types of SCFAs. Overall, the overrepresentation and extensive distribution of several CAZyme domains in copresent rumen MAGs *vs.* all MAGs indicated that the copresent rumen MAGs played a crucial role in rumen and *in vitro* fermentation for degrading plant biomass.

### 3.6 Relationships between copresent rumen MAGs and short-chain fatty acids production in RUSITEC

The results of Pearson analysis showed significant correlations (r > 0.75, Q- value < 0.05) between abundance of copresent rumen MAGs and specific SCFA concentration in RUSITEC. The 14 MAGs with both genome-level evidence of producing SCFAs and positive correlations with SCFAs concentration were obtained for further confirmation based on the gene of producing SCFAs in MAGs. Among these 14 MAGs, 10 MAGs and 3 MAGs were associated with acetic acid and propionic acid production, respectively, while 1 MAG was associated with the production of butyric acid and propionic acid. Moreover, there were multiple positive correlations between these copresent MAGs. From the above results it is revealed that the SCFAs were co-produced by several positively correlated copresent rumen MAGs.

The MAGs associated to the acetic acid, propionic acid and butyric acid production were classified into 8 different bacterial families. There were 3 of 14 MAGs were classified into species level. Simultaneously, 2 MAGs (C6_17, C7_53), which were classified into *Ruminococcus*, were correlated with producing propionic acid, and acetic acid in RUSITEC, and these two copresent rumen MAGs were annotated to specie level of *Ruminococcus_E sp900320415* and R*uminococcus_E sp900100595*, respectively. The remaining 1 species-levels MAG (M10_35: family *Lachnospiraceae*; genus *UBA629*; species *900317915*) was simultaneously associated to the propionic acid and butyric acid production in RUSITEC. The higher-level taxa to which these MAGs belong have been shown in previous studies to be associated with the production of SCFAs ^39–40^. On the other hand, it was also found that members of rumen *Lachnospiriaceae* belonged to heritable rumen species ^41^. With the addition of rumen *in vitro* fermentation conditions in current study, some members of *Lachnospiriaceae* family were still the copresent rumen microbiome.

For MAGs classified into unknown species, these MAGs were classified into different genera (genus *Lachnobacterium*: M11_55; *Prevotella*: C6_59; *RC9*: C7_26; *RUG472*: G4_308; RUG740: M9_208; *UBA1777*: C5_192, C6_77, C7_25, C7_99, OC1_56; *UBA2866*: M9_188). *Ruminococcus*, *Prevotella*, and *RC9* genera were the main microbial taxa in rumen, according to earlier research, and these genera were linked to the concentration of SCFAs in distinct digestive tracts ^42–45^. Despite the fact that some of these MAGs and genus have not been studied recently, the correlation analysis in this work and the linked genes derived from MAGs may be utilized as evidence for synthesizing SCFAs in the rumen and RUSITEC. Moreover, some previously unclassified taxa that are related to the SCFAs production during *in vitro* ruminal fermentation were observed, adding to our knowledge of rumen microbial activity under *in vitro* ruminal fermentation.

In conclusion, the different microbial communities and SCFAs production in RUSITEC were driven by the initial inoculum. Nonetheless, of all 1677 recovered MAGs, the copresent rumen microbiome consisting of 298 MAGs overrepresented in functions of degrading xylose/pectin and specific pathways, could still exist in rumen and RUSITEC samples, as well as other different metagenomic dataset collected from previous study. Additionally, some specific and unannotated copresent rumen MAGs associated with the SCFAs production in RUSITEC were observed. This study provides new insights into the copresent rumen microbiome which could be considered as candidate key scopes for research on *in vitro* ruminal fermentation.

## Abbreviations Used

SCFA: short-chain fatty acids
RUSITEC: rumen simulation technique
MAG: metagenome-assembled genome
CAZymes: Carbohydrate-Active enZYmes
KEGG: Kyoto encyclopedia of genes and genomes
ANI: average nucleotide identity
PCoA: principal co-ordinates analysis
PL: polysaccharide lyase
GH: glycoside hydrolase
CBM: carbohydrate-binding module
SLH: S-layer homology module
AA: auxiliary activities
CNGB: China National GeneBank.

## Supporting Information

Classification and completeness of 298 copresent MAGs (Table S1) (XLS) Archaeal phylogenetic tree and the overrepresentations of KEGG pathway in copresent MAGs vs all MAGs (Figures S1–S3) (DOC)

## Acknowledgement

This work was supported by National Natural Science Foundation of China (31822052); and China National GeneBank (CNGB).

## Data Availability

The data underlying this study are openly available in China National GeneBank at https://db.cngb.org/cnsa/project/CNP0002970_af2e0480/reviewlink/.

