## Supplementary data for "Copresent microbiome and short-chain fatty acids profiles of plant biomass utilization in rumen simulation technique system"

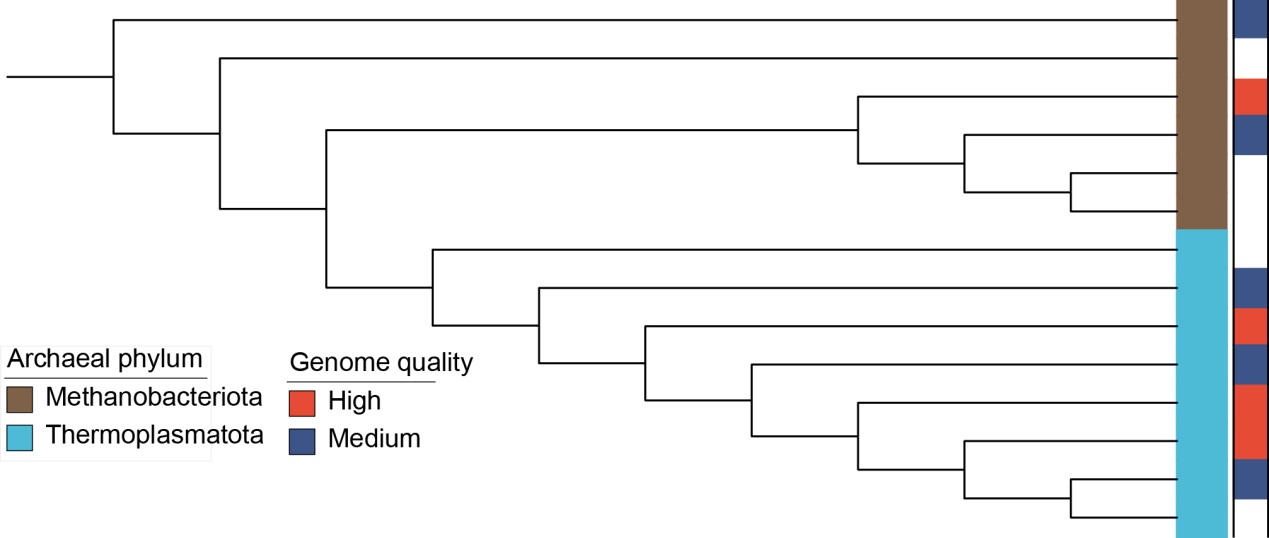

Figure S1. Taxonomic tree of archaea based on 122 single copy genes. Color of right cell corresponds to the genome quality of each archaeal MAGs; the color of left cell represents the different phylum of archaeal MAGs.

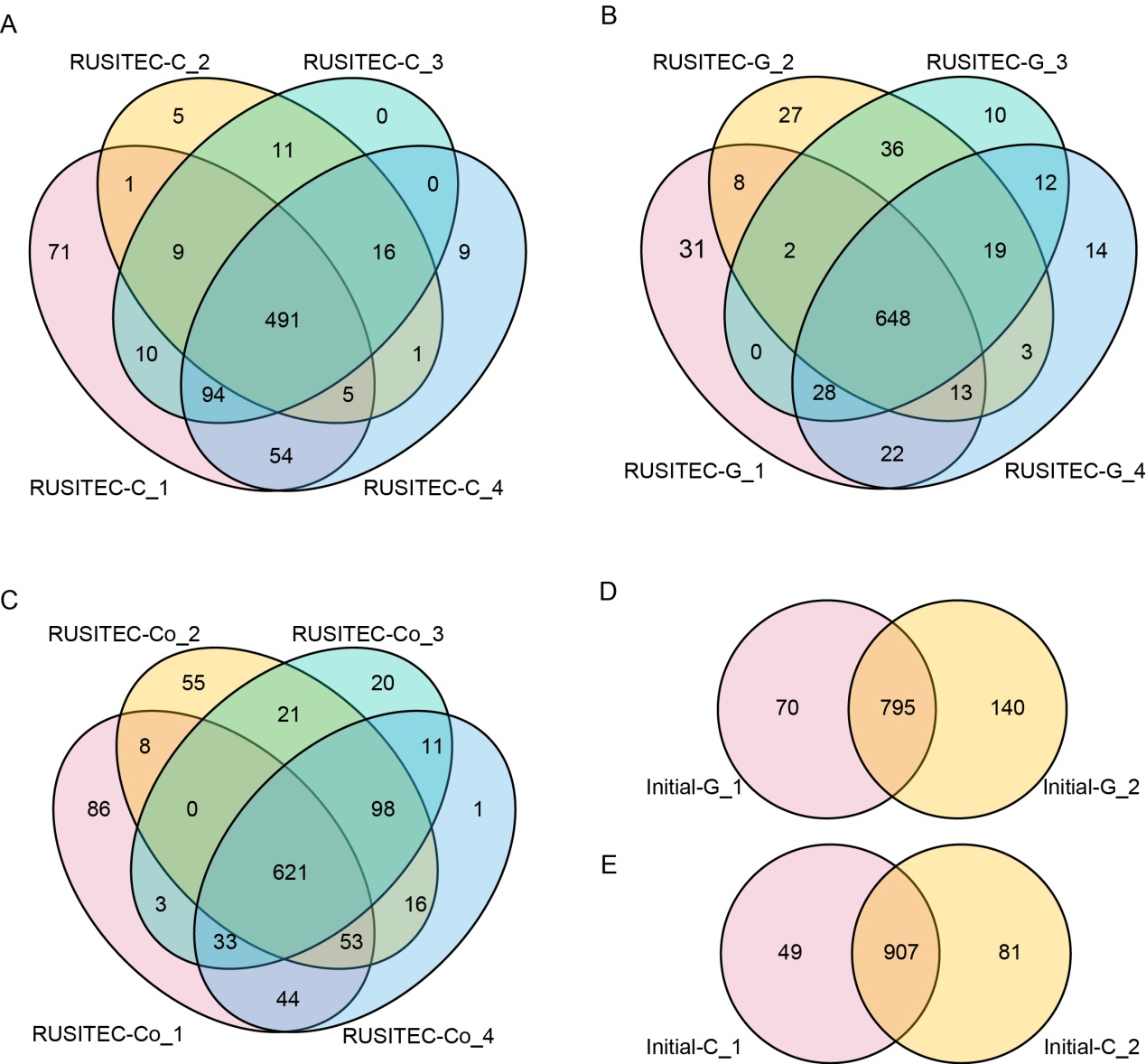

Figure S2. A-E: Venn diagram shows the presence of MAGs across every sample in each group based on MAG mapping method; The RUSITEC-C_1-4 and RUSITEC-G_1-4, respectively represent RUSITEC inoculated with rumen contents from cow and goat, RUSITEC-Co_1-4 represent RUSITEC inoculated with the biomass-normalized mixtures of rumen contents from cow and goat, initial_G_1-2 and initial-C_1-2 represent rumen contents collected from cow and goat respectively.

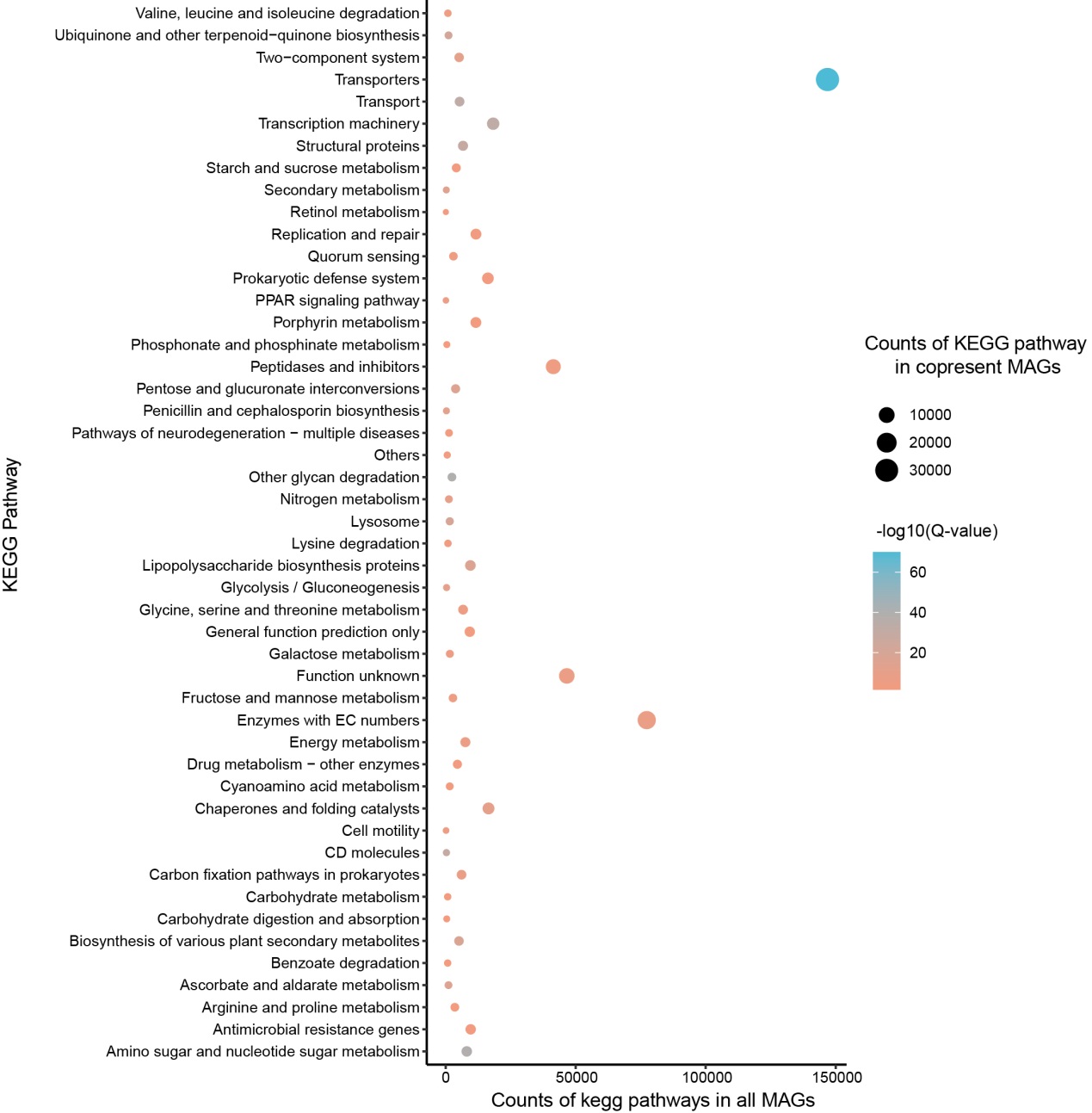

Figure S3. The overrepresentations of KEGG pathway in copresent MAGs vs all MAGs. KEGG pathway counts are shown in log-scale. The size of points indicates the count of KEGG pathway in copresent rumen MAGs, and the color of points indicates –log10 (Q-value) of Fisher's exact test.
